# *Lef1* is dispensable for blood-brain barrier integrity despite its dominant role in endothelial Wnt signaling

**DOI:** 10.64898/2026.03.14.711770

**Authors:** Yoel Yeretz-Peretz, Shira Anzi, Batia Bell, Ayal Ben-Zvi

## Abstract

The blood-brain barrier (BBB) is a specialized vascular structure essential for CNS homeostasis, whose formation and maintenance are governed by the canonical Wnt signaling pathway. While the upstream ligands and receptors are well-characterized, the downstream transcriptional architecture remains poorly understood. Here, we investigate the functional requirement for Lef1, the most abundant Tcf/Lef transcription factor and a hallmark readout of Wnt activity in brain endothelial cells (BECs). Utilizing a conditional, endothelial-specific deletion strategy in mice, we demonstrate that loss of *Lef1* during either embryonic development or adult homeostasis significantly dampens Wnt transcriptional output. Surprisingly, high-resolution molecular and functional analysis reveals that this reduction results in only minor dysregulation of the BBB-specific gene program and is insufficient to trigger barrier breakdown. Our results establish that Lef1 is not an obligatory effector of the Wnt-dependent BBB maintenance program. These findings suggest a resilient transcriptional framework where redundant Tcf/Lef family members or alternative regulatory circuits preserve CNS microvasculature function, providing new insights into the genetic robustness of the blood-brain barrier.

**Research Highlights:**

- **Lef1 is the predominant Tcf/Lef transcription factor** in the brain endothelium and a hallmark of vascular Wnt signaling.
- **Endothelial-specific *Lef1* deletion** significantly attenuates Wnt transcriptional output during both CNS vascular development and adult homeostasis.
- **Loss of Lef1 causes only minor perturbations** in the BBB-specific gene program, rather than a global loss of endothelial identity.
- **Structural and functional BBB integrity is preserved** in the absence of Lef1, revealing a high degree of vascular resilience.
- **The findings demonstrate that Lef1 is not an obligatory effector** for the maintenance of the specialized CNS microvasculature.

## 1. Introduction

Brain endothelial cells (BECs) composing most of the capillaries in the central nervous system (CNS) form a highly selective barrier that controls the influx and efflux of ions, nutrients, metabolites, signaling molecules, and immune cells from the blood into the brain parenchyma and vice versa. This unique structure, termed the blood-brain barrier (BBB), is composed also of pericytes and astrocytic end-feet interacting with the endothelial cells and thus allows maintenance of a homeostatic environment in the CNS, critical for proper neuronal function^1^. Breakdown of BBB functionality has been strongly linked to neurological affliction and dysfunction such as neuro-degenerative, neuro-inflammatory diseases, trauma and more^2–8^.

The selective nature of the BBB limits paracellular and transcellular movement of ions, molecules, and cells through the layer of endothelial cells, as well as actively facilitating transport of required components between the blood and the parenchyma. Paracellular diffusion is restricted by the presence of tight junction proteins like claudin 5 (referred hereafter as *cldn5*) in addition to the adhesion molecules connecting neighboring endothelial cells^9^. Transcellular movement of ions and molecules is restricted by the complete absence of fenestra and significantly reduced vesicular transport (transcytosis), particularly CAV1-mediated transport. Transcytosis restriction is thought to be the result of high *Mfsd2a* expression, a membrane lipid flippase that modifies endothelial membrane lipid composition, thereby limiting vesicle formation^1,10,11^. In addition, the BECs also express various selective transporters such as the *Glut1* glucose transporter, as well as active ATP dependent ABC transporters, also termed multidrug resistance transporters, such as p-glicoprotein (P-gp, known also as MDR1, or ABCB1) and the breast cancer resistance protein (BCRP known also as ABCG2)^1^.

Canonical Wnt signaling pathway is crucial in regulating BBB development from angiogenesis and brain vascularization to BBB formation^12–19^, as well as being critical for BBB maintenance in adulthood^20,21^. Wnt7a/b and norrin ligands secreted from embryonic neural progenitor cells and postnatal astrocytes^18,22–24^ have been shown to bind BECs and signal through a specific sub-set of receptors and co-receptors^13,14,16,17,25^. Signaling leads to stabilization of β-catenin, facilitating its entry into the nucleus where together with members of the *Tcf/Lef* transcription factor family it induces transcription of pro BBB genes (e.g. *Mfsd2a*, *Glut1*, and *Cldn5*) and suppresses the expression of genes characteristic of non-barrier capillaries such as the fenestra structure component *Plvap*^19–21,26,27^.

Extensive research has established the critical roles of Wnt ligands and receptors^13,14,16–18,22,25^, and of β-catenin^15,19–21,28^ in BBB development and maintenance. In contrast, the *Tcf/Lef* contribution is only implied, by extrapolating from other biological systems in which this pathway is active^29^, and by an enrichment of accessible *Lef1* and *Tcf7* binding motifs in BECs compared to peripheral endothelium^26,30,31^. Among the four *Tcf/Lef* genes expressed in vertebrates, namely *Tcf7* (*TCF1*), *Lef1*, *Tcf7l1* (*TCF3*), and *Tcf7l2* (*TCF4*)^29^, *Lef1* is the most commonly associated with Wnt signaling in the BBB evident by an expression of over 40 fold mRNA levels in the brain vs. lung capillaries, compared to *Tcf7* (*Lef1* has 144 times more average sequencing reads in the brain than in the lung, while *Tcf7* has 3.24 times more)^32–34^. *Lef1* being a target gene by itself, due to a positive feedback regulation, is also commonly used to validate loss of Wnt signaling in knock-out (KO) models of Wnt ligands, receptors, and β-catenin^21,26^.

In the present study we aim to further elucidate the role of *Tcf/Lef* in BBB Wnt signaling. We tested if *Lef1* is necessary for Wnt signaling in BECs or if other *Tcf* are compensating in its absence. To this end, we utilized transgenic mice with floxed *Lef1* alleles and endothelial-specific Cre expression to achieve *Lef1*-KO either induced in adulthood or during embryonic development and tested the effect on BBB integrity. While we could validate *Lef1* loss, and detect some minor changes in the expression of Wnt target BBB genes, we did not observe any significant functional defects in BBB functionality, pointing to a likely compensation from other *Tcf/Lef* family members.

## 2. Results

### 2.1. Use of endothelial tropic AAVs to induce *Lef1* loss of function mutagenesis in adult mice

Utilizing published single cell RNAseq data (‘Betsholzlab’ resource)^32–34^, we compared *Tcf/Lef* expression in brain and lung endothelial cells. This comparison revealed *Lef1* to be the most abundant *Tcf/Lef* gene in BECs, with a mean expression of 365 reads in brain endothelium compared to a mean 2.53 reads in lung endothelium. The only other *Tcf* family member showing a significant increased expression in BECs is *Tcf7*, with a mean 38.5 reads in brain endothelium and 11.9 mean reads in lung endothelium. No significant difference in expression could be detected in BECs and lung endothelium for reads of *Tcf7l1* (mean 4.19 and 2.97 reads respectively) and of *Tcf7l2* (mean 23.0 and 21.8 reads respectively) (Figure 1A). Based on the relative high expression and on the extremally robust deferential expression in BBB vs non-BBB ECs (144-fold higher), our working hypothesis was that Lef1 might play a critical role in the reported β-catenin dependent barrier maintenance function at the BBB^20,21^.

**Figure 1.**
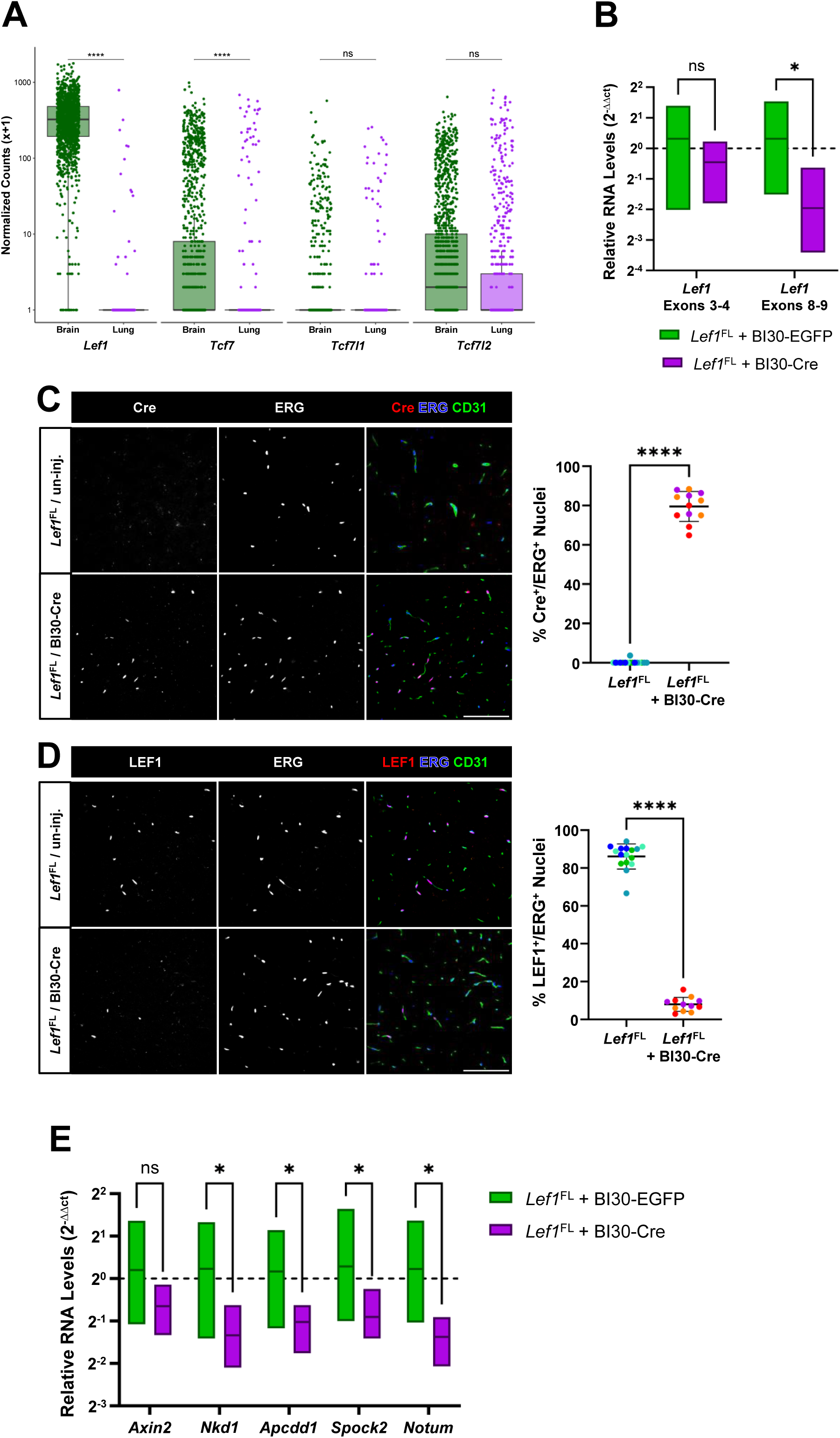
Use of endothelial tropic AAVs to induce *Lef1* loss of function mutagenesis in adult mice. **(A)** Box plots of *Tcf/Lef* expression levels in single-cell RNAseq data of vasculature, taken from mice brain and lung (analysis of the ‘Betsholtzlab’ data^32–34^). Normalized counts +1 presented at log10 scaling (multi-comparison t-test was performed on raw data before x+1 transformation). **(B)** Relative isolated capillaries mRNA levels (qPCR) of *Lef1* show no changes in total *Lef1* transcripts (Exons 3-4), while a significant reduction is found in the wild-type *Lef1* transcripts (Exons 8-9). mRNA levels from *Lef1*-KO capillaries (*Lef1*^FL^ + BI30-Cre) are compared to control mice (*Lef1*^FL^ + BI30-EGFP). **(C)** Efficient and specific viral infection, drives Cre expression. Representative epi-fluorescent images of immunostained Cre^+^ brain endothelial cells in the cortex of AAV-BI30-Cre injected *Lef1*^FL^ mice and un-injected controls. Quantification shows 79.33% increase of Cre^+^ endothelial nuclei in cerebral cortex of injected mice (anti-CD31 used to mark endothelium and anti-ERG to mark endothelial nuclei). **(D)** Quantification of LEF1 staining reveals a 90.73% decrease in LEF1^+^ endothelial nuclei (co labeled with anti-ERG). For microscopy n=4 fields of n=4 control and n=3 *Lef1*-KO mice were used, data is presented as mean and standard deviation, colors represent individual mice, two-tailed Welch’s t test was used, ****p < 0.0001, scale bars are 100µm. **(E)** Relative mRNA levels (qPCR) of known Wnt signaling target genes in isolated capillaries, show significant reduction in *Nkd1*, *Apcdd1*, *Spock2*, and *Notum* following BI30-Cre injection. qPCR was performed on n=6 samples from each treatment, data is represented as the relative expression normalized to the average expression in control samples transformed to 2^-ΔΔCt^, two-tailed Welch’s t test was used.

Next, we created endothelial specific *Lef1*-KO mice (loss of function mutagenesis) by introducing AAV-BI30 viral vector driving expression of Cre-recombinase (henceforth BI30-Cre) into adult B6.Cg-*Lef1^tm1Hhx^*/J (henceforth *Lef1*^FL^)^35^ mice 3-4 weeks prior to sacrificing the animals (Supplementary figure 1A). In these mice, exon 7 and 8, which encode *Lef1* DNA binding domain, are flanked by LoxP sites allowing to render *Lef1* ineffective by using Cre^35^. For Cre delivery we utilized AAV-BI30, which is highly efficient in transducing endothelial cells in the CNS of different mice stains^36^, resulting in close to 80% Cre^+^ BECs (79.56 ± 7.61% in the *Lef1*-KO vs. 0.23 ± 0.93% in the un-injected controls, all results presented as mean ± standard-deviation) with no significant nuclear staining of Cre outside of ERG^+^ endothelial cells (Figure 1C and supplementary figure 1B).

As BI30 is reported to be strongly biased towards transducing BECs but still transduces some peripheral endothelial cells, for practical reasons we genotyped DNA from lung samples of *Lef1*^FL^ mice injected with BI30-Cre or BI30-EGFP (control) revealing excision of exons 7-8 only in BI30-Cre injected animals (Supplementary figure 1C). To test the efficiency of genetic perturbation at the mRNA level, we performed quantitative real time PCR (qPCR) and evaluated relative levels of mRNA in isolated brain capillaries. By using primers recognizing *Lef1* exons 8 and 9 (beyond the targeted floxed region) we observed about 5 times reduction of *Lef1* mRNA (Figure 1B). In contrast, when using primers that recognize *Lef1* exons 3 and 4 (before the targeted floxed region), no significant reduction in mRNA could be observed (Figure 1B). This result indicates that excision of exons 7-8 probably does not impact *Lef1* transcription, and that a truncated *Lef1* transcript is probably expressed in the KO cells at similar level as the WT *Lef1* transcript in the control mice.

We then tested if the reduction in mRNA does also lead to a decrease in LEF1 protein levels in BECs. To that end we co-stained brain slices from *Lef1*-KO and control mice for LEF1 and for the endothelial specific transcription factor ERG. While 86.13 ± 6.65% of ERG^+^ BECs were also LEF1^+^ in the cortices of control mice, *Lef1*-KO mice showed only 7.98 ± 3.68% LEF1^+^ BECs, a reduction of ∼90% (Figure 1D). Thus, percentage of LEF1 suppression aligns well with percentage of Cre expression, and slight discrepancy between them can be attributed to differences in antibody staining efficiency. Overall, these results confirm that the usage of BI30-Cre in *Lef1*^FL^ mice, can consistently lead to broad reduction of LEF1 in BECs making it an appropriate tool for testing the effects of *Lef1* loss of activity on BBB function.

Contrary to the reported neurological pathologies that result from *Ctnnb1*-KO 1-2 weeks following KO induction^21^, no such effects were observed in our *Lef1*-KO even 4 weeks after AAV-BI30 injection. We therefore tested if loss of *Lef1* in adult mice impacts canonical Wnt signaling similarly to loss of β-catenin. We evaluated mRNA levels of several known Wnt target genes previously shown to be downregulated in models of endothelial specific β-catenin-KO^20^. *Nkd1, Apsdd1, Spock2,* and *Notum* were significantly reduced by about a factor of 2, and *Axin2* exhibited a decrease trend but was not statically significant (Figure 1E). Together with the absents of a significant change in *Lef1* expression (itself also being a down-stream target of Wnt signaling) (Figure 1B), this result suggests that *Lef1* is mediating part of Wnt signaling effects (compared to β-catenin) as indicated by the partial overlapping effect of *Ctnnb1*-KO on the down-stream target expression^20^.

Overall, these results show that our approach results in extensive *Lef1*-KO specifically in the BECs without significantly infecting other cell types in the brain, while also affecting some of Wnt signaling. This will allow us to reliably test the loss of *Lef1* effect in BECs, on maintenance of BBB function.

### 2.2. BECs *Lef1* loss of function has no detectable effect on canonical Wnt signaling induced expression of BBB related downstream targets

In order to evaluate the potential effect on the BBB, we tested if the loss of *Lef1* in BECs leads to changes in BBB specific genes expression, previously shown to be regulated by Wnt signaling. Along the study we used CD31 (PECAM1), isolectin-IB_4_, and Laminin known as general vasculature markers, and tested co-immunostaining with barrier and peripheral EC markers^37^. First, we assessed the expression of *Plvap* co-immunostaining in cerebral cortices of *Lef1*-KO and control mice (un-injected). *Plvap* is a component of fenestra and is therefore absent from most of the brain blood-vessels. Previous research on the role of canonical Wnt signaling in postnatal mice, in glioma, and in the subfornical organ established a strong connection between Wnt signaling and downregulation of *Plvap*^19,38,39^. We therefore expected that if *Lef1* has a critical role in the Wnt signaling pathway of BECs, *Lef1*-KO will lead to *Plvap* upregulation. Surprisingly no such upregulation could be observed in the absences of *Lef1* (Figure 2A). To ensure that this was not due to a technical immunostaining issue we also quantified PLVAP^+^ blood vessels in the choroid plexus (henceforth ChP), which is known to be fenestrated^40^, and observed robust coverage as expected, with no significant change between the control and *Lef1*-KO mice (Supplementary figure 2A). This result was further validated by measuring *Plvap* mRNA levels in cortical isolated capillaries, resulting in no significant changes between the groups (Figure 2D).

**Figure 2.**
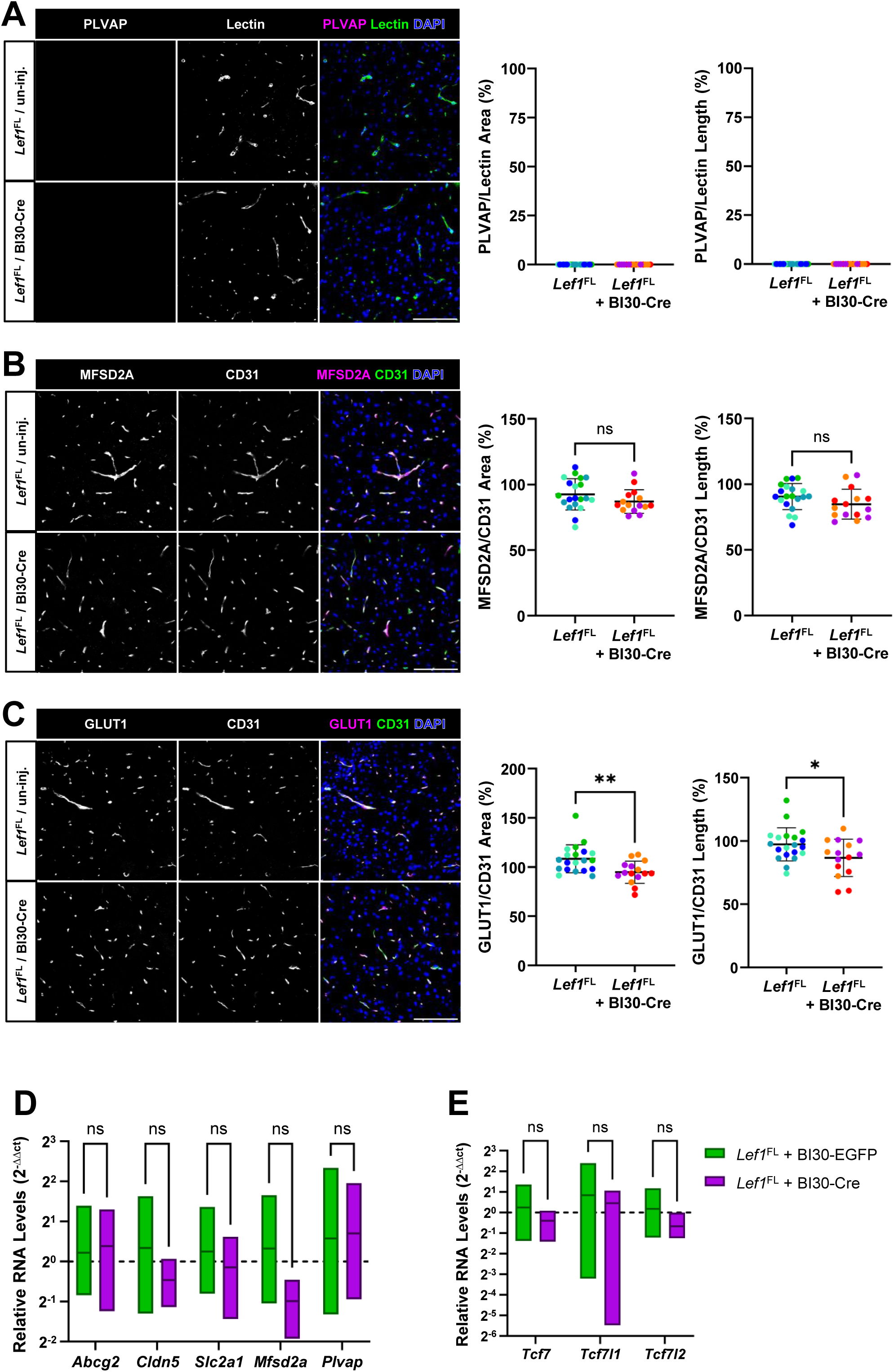
BECs *Lef1* loss of function has no detectable effect on canonical Wnt signaling induced expression of BBB related downstream targets. **(A)** Representative confocal images of PLVAP^+^, **(B)** MFSD2A^+^, **(C)** and GLUT1^+^ endothelial cells immunostaining (anti-CD31 or Isolectin) in the cortex of AAV-BI30-Cre injected *Lef1*^FL^-KO mice and un-injected controls. Quantification of n=5 fields taken from n=4 control and n=3 *Lef1*-KO mice, reveal a statistically significant decrease in coverage, only in the case of GLUT1 (12.59% and 10.89% in the area and length respectively) as a result of *Lef1*-KO. Data is presented as mean and standard deviation, different colors represent individual mice, two-tailed Welch’s t test was used, ns = non-significant, *p < 0.05, **p < 0.01, scale bars are 100µm. **(D)** Relative mRNA levels (qPCR) of BBB genes related to Wnt signaling regulation **(E)** and of the other *Tcf* family members in isolated capillaries from *Lef1*-KO (*Lef1*^FL^ + BI30-Cre) and control mice (*Lef1*^FL^ + BI30-EGFP). This analyses show no significant changes in the expression of any of the tested genes. qPCR was performed on n=6 samples from each treatment, data is represented as the relative expression normalized to the average expression in control samples transformed to 2^-ΔΔCt^, two-tailed Welch’s t test was used.

While some previous reports indicated that BECs specific KO of *Ctnnb1* not necessarily resulted in *Plvap* upregulation^20^, many BBB specific genes have previously been shown to be regulated by the canonical Wnt signaling pathway. In order to determine whether *Lef1* deletion affects the expression of these genes, we performed immunostaining for four established blood–brain barrier markers; glucose transporter 1 (GLUT1)^26^, sodium-dependent lysophosphatidylcholine symporter 1 (MFSD2A)^20^, tight junction protein claudin 5 (CLDN5)^20^, and breast cancer resistance protein (BCRP, also known as ABCG2)^27,41^. In concordance with our PLVAP results, no significant changes in MFSD2A (Figure 2B), CLDN5 (Supplementary figure 2B) and BCRP (Supplementary figure 2C) were observed. The only modest but statistically significant change was observed with immunostaining for GLUT1, resulting in GLUT1 vascular coverage decreasing from 108.40 ± 14.20% blood-vessel area and 97.28 ± 13.06% vessel length in the control to 94.75 ± 11.28% area and 86.63 ± 14.72% length in *Lef1*-KO, a 11-13% decrease (Figure 2C). Higher than 100% blood-vessel coverage (area or length) is possible since the values are normalized to CD31 or Isolectin staining which might be less efficient than some more efficient antibodies like anti-GLUT1.

Performing qPCR for those four genes on mRNA of isolated brain capillaries, indicated no statistically significant reduction. Reduction trends for *Glut1*, *Mfsd2a* and *Cldn5* mRNA levels were noted but were not statistically significant (Figure 2D). Taken together those results indicate that *Lef1* is not necessary for maintaining expression of BBB specific genes.

As mentioned, in homeostatic conditions the steady-state level of *Lef1* is the most abundant of the *Tcf* family members. Nevertheless, based on our results we hypothesized that one or more of the other *Tcf* family members may be upregulated to compensate for the loss of *Lef1*. Thus, we performed qPCR on the same isolated brain capillaries as before, to test the relative levels of the other three *Tcf* genes. At this level of regulation, we could not detect significant changes in any of the *Tcf* genes comparing between the control and *Lef1*-KO mice (Figure 2E). Of note, capillary isolation enriches all vasculature cell types, and since *Lef1*-KO is endothelial specific, only strong changes in gene expression are expected to be captured with this method. Thus, it is possible that some level of compensatory upregulation is not detected. In addition, *Tcf7* is already upregulated in the BECs compared to lung endothelial cells, although not to the level of *Lef1* (Figure 1A), and might be able to compensate for the lack of *Lef1* at its steady-state level. Therefore, further research is needed in order to elucidate the possible role of *Tcf7* in BBB maintenance.

Altogether, *Lef1*-KO seemed to have no significant effect on the expression of BBB associated genes known to be regulated by canonical Wnt signaling, with GLUT1 being the only exception with modest decrease.

### 2.3. Loss of *Lef1* function in BECs has no observable effect on the expression of downstream Wnt signaling targets in susceptible brain areas

Recent, pre-print study findings show that loss of canonical Wnt signaling via *Ctnnb1*-KO in BECs of adult mice causes damage mainly in the cerebellum, specifically at the molecular layer (CML) and hippocampus, and less so in the cortex^21^. This reflects earlier results in post-natal mice, where BEC specific KO of various genes of the canonical Wnt signaling pathway resulted in a much more dominant effect in the cerebellum than in the cerebral cortex^42^. We therefore expanded our analysis to the cerebellum to test if any significant changes in the previously mentioned downstream targets could be observed in the CML. To that end we co-immunostained for the downstream targets and CD31/isolectin-IB_4_ as a reference marker and quantified the vascular area and length covered by the target marker. In this analysis we excluded vessels in the granular layer, as well as the arteries in the CML in the case of MFSD2A (red areas in figure 5 and supplementary figure 5 are marking the areas excluded from the analysis), due to their naturally low *Mfsd2a* expression in arteries^33^.

In concordance with our cortex results we found that in the cerebellum there were no significant changes in the coverage of PLVAP (Figure 3A), GLUT1 (Figure 3B), and BCRP (Supplementary figure 3A). Although significant changes were observed in MFSD2A and CLDN5. MFSD2A^+^ vessel length decreased by 12.15% in the cerebellum of *Lef1*-KO mice (from 97.71 ± 22.06% in control mice to 85.84 ± 11.17% in KO mice), but MFSD2A^+^ vessel area was not significantly impacted (Figure 3C). In the case of CLDN5^+^, an opposite effect of a minor but statistically significant increase of 8.34% in vessel area could be observed in the cerebellum of *Lef1*-KO mice (from 78.88 ± 8.22% in control mice to 85.46 ± 10.01% in KO mice), and vessel length showed an increase of 8.94% in KO mice that was not statistically significant (Supplementary figure 3B).

**Figure 3.**
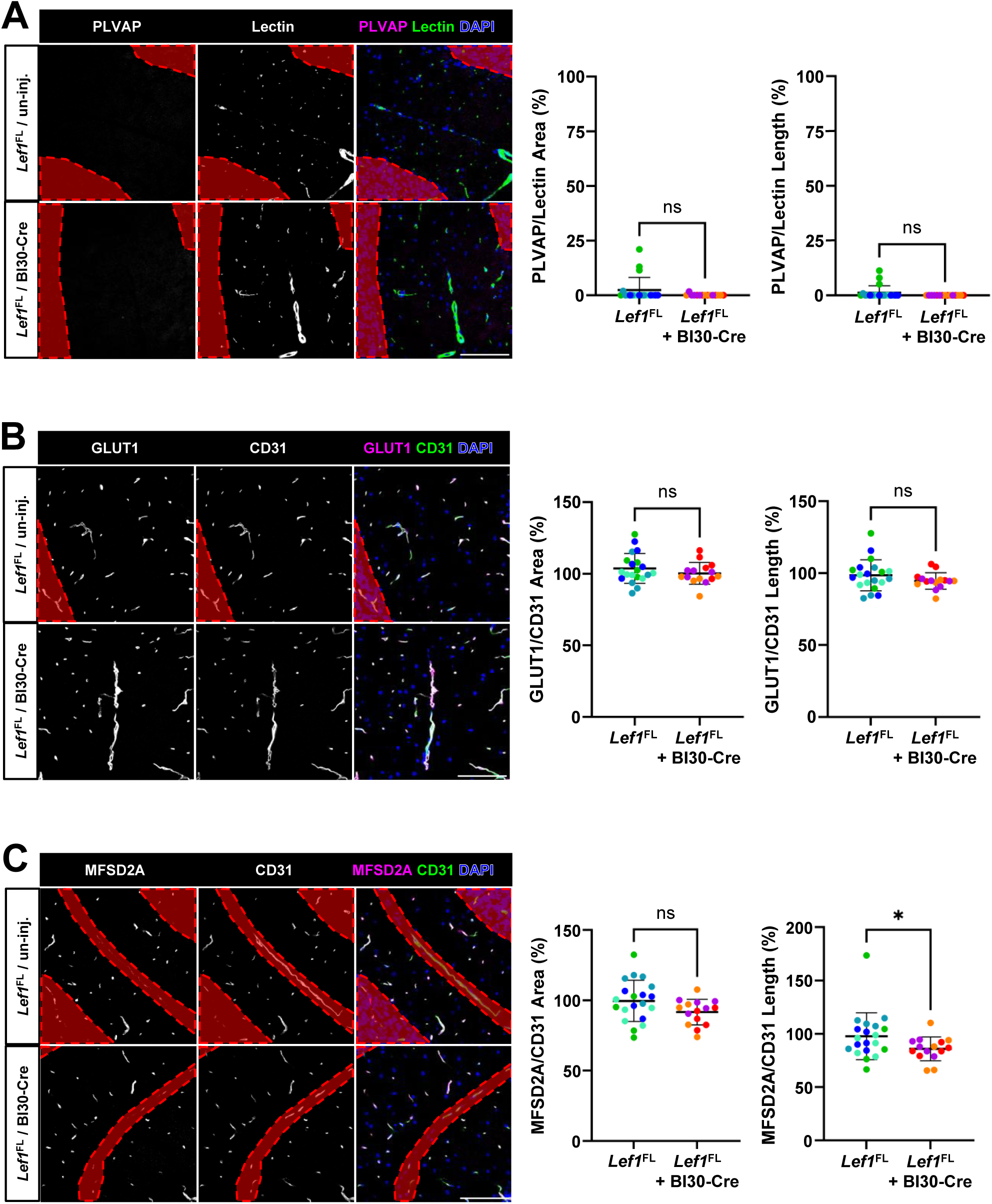
Loss of *Lef1* function in BECs has no observable effect on the expression of downstream Wnt signaling targets in susceptible brain areas. **(A)** Representative confocal images of PLVAP^+^ **(B)** GLUT1^+^, **(C)** and MFSD2A^+^ endothelial cells immunostaining (anti-CD31 or Isolectin) in the cerebral molecular layer of AAV-BI30-Cre injected *Lef1*^FL^-KO mice and un-injected controls. Quantification of n=5 fields taken from n=4 control and n=3 *Lef1*-KO mice, reveal a statistically significant decrease in coverage only in the case of GLUT1 (12.15% in the length) as a result of *Lef1*-KO, when the analyzed area is restricted to only the molecular layer and excludes the arteries in the case of MFSD2A (non-molecular layer areas in red were excluded from analyses). Data is presented as mean and standard deviation, different colors represent individual mice, two-tailed Welch’s t test was used, ns = non-significant, *p < 0.05, scale bars are 100µm.

In total, loss of *Lef1* seems not to replicate effects related to loss of *Ctnnb1* on expression of BBB associated genes in the cerebellum either. The modest changes observed in MFSD2A and CLDN5 are inconsistent in their direction.

### 2.4. Loss of *Lef1* function in BECs has no effects on BBB permeability

Loss of Wnt signaling in BECs, either by knocking out the Wnt receptors or β-catenin, has been shown to result in a leaky barrier in postnatal^19,42^ and adult^20,21^ mice. Although *Lef1*-KO in BECs of adult mice did not result in the expected effect on gene expression, other mechanisms might be affecting BBB function, and so we turned to examine barrier integrity in this model. To do so, we perfused BI30-Cre injected *Lef1*-KO mice and un-injected control mice with sulfo-NHS-biotin, a ∼500 Dalton tracer previously shown to leak in other Wnt signaling impeded models^21,42^. No biotin leakage could be detected in either the cortex, the thalamus, or the cerebellum of *Lef1*-KO mice, despite the marked decrease in LEF1^+^ BECs (Figure 4A). This contrasted with the pronounce biotin leakage that could be observed in the ChP which naturally contains leaky non-barrier blood vessels ^40^ (Supplementary figure 4A).

**Figure 4.**
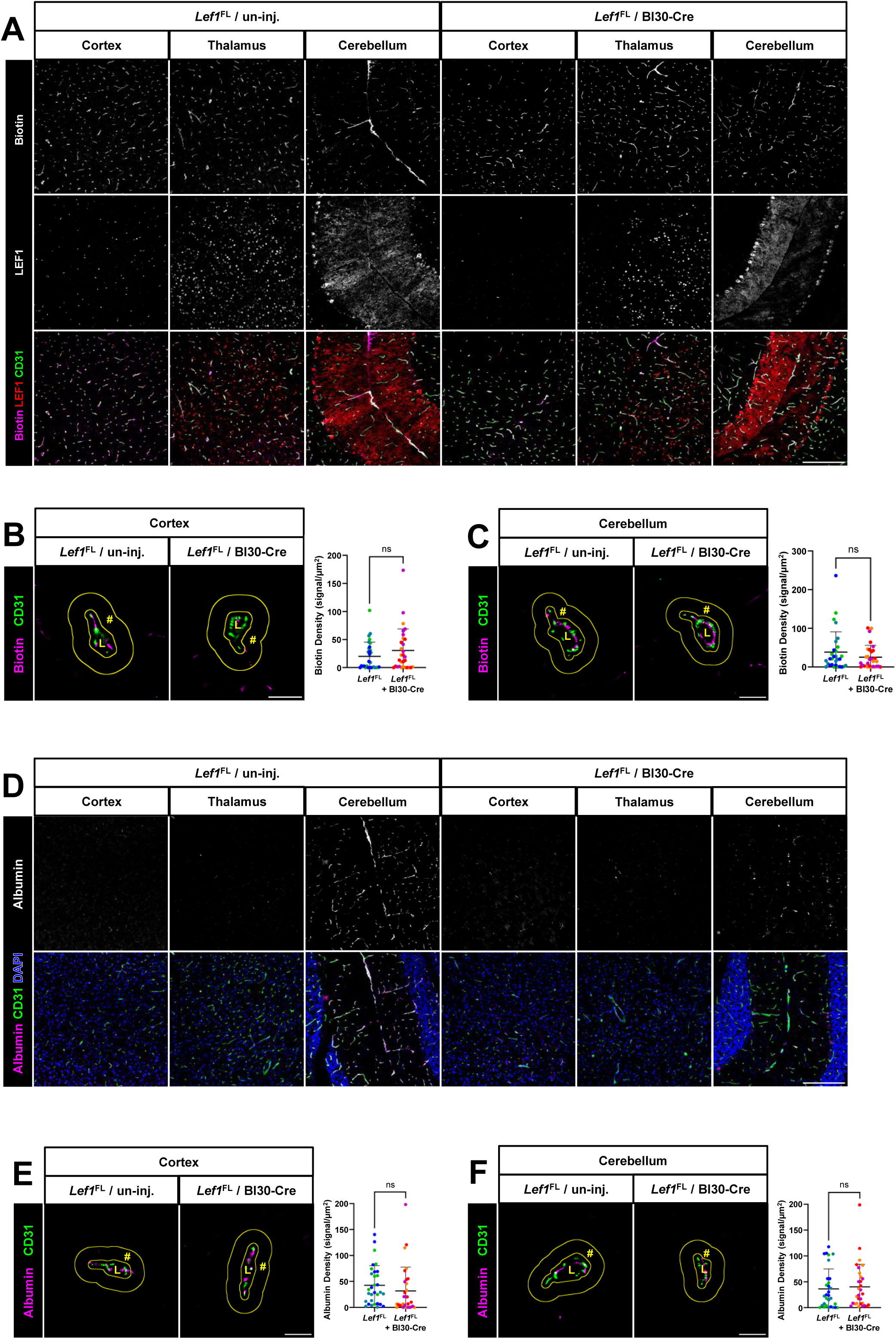
Loss of *Lef1* function in BECs has no effects on BBB permeability. **(A)** Representative confocal images of sulfo-biotin tracer challenges in the cortex, thalamus, and cerebellum of AAV-BI30-Cre injected *Lef1*^FL^ mice and un-injected controls. No detectable biotin leakage was apparent in any of the three areas. Scale bar is 200µm. **(B)** Quantification of super-resolution microscopy images (dSTORM) of capillaries cross sections, from the cortex and **(C)** the cerebellum of the same mice confirmed the lack of biotin leakage induced by the loss of *Lef1* function. L = lumen, # = area abluminal to the endothelium quantified, scale bar is 2µm. **(D)** Similarly, representative confocal images of endogenous albumin immunostaining in the cortex, thalamus, and cerebellum do not show any leakage neither. Scale bar is 200µm. **(E)** Quantification of super-resolution images (dSTORM) of capillaries from the cortex and **(F)** the cerebellum confirms the lack of albumin leakage induced by the loss of *Lef1*. L = lumen, # = area abluminal to the endothelium quantified, scale bar is 2µm. Quantification of super-resolution data was done on n=10 capillaries cross profiles, of n=3 mice of each treatment, data is presented as mean and standard deviation, different colors represent individual mice, two-tailed Welch’s t test was used, ns = non-significant.

To quantify biotin leakage across the BBB with higher sensitivity, we utilized Stochastic Optical Reconstruction Microscopy (dSTORM). This super-resolution approach allows us to localize biotin on a nanometer scale. Moreover, it allows high level of sensitivity, being a single molecule microscopy approach (SMLM), in case we miss delicate changes in BBB function using confocal microscopy. With this technique, we quantified the number of biotin molecules that crossed the endothelial cell layer, which we defined as 250 nanometers ab-luminal to the CD31 marker, and are up to 1µm ab-luminal. This high-resolution quantification reinforced the observation that *Lef1*-KO does not affect BBB permeability to small tracers, neither in the cortex (Figure 4B) nor in the CML (Figure 4C). Quantifying leakage relative to the astrocytic marker AQP4, similarly showed no enhanced leakage into the brain parenchyma of the cortex in the absence of *Lef1* expression (Supplementary figure 4B).

We also tested for leakage of an endogenous tracer, namely albumin which is also known to be leaking in BEC specific *Ctnnb1*-KO models^20^. For this analysis we used the same biotin perfused samples, this resulted in the removal of most vascular albumin but is not expected to affect albumin that extravasated the blood vessels. Similar to biotin, no increased leakage could be detected either in the cortex or CML (Figure 4D), while non-barrier blood vessels of the ChP were as expected, hyper permeable (Supplementary figure 4C). dSTORM analysis of albumin relative to CD31 in the cortex or CML (Figure 4E and 4F), and relative to AQP4 (Supplementary figure 4D) confirmed that no increased albumin leakage occurs in the absences of endothelial *Lef1* expression.

As expected from the lack of significant changes in the expression of BBB associated genes following *Lef1*-KO in adult mice, the loss of *Lef1* also failed to impact BBB integrity when challenged with a small sized exogenous tracer, as well as for a larger endogenous tracer. This strengthens our conclusion that *Lef1* is not necessary for maintaining Wnt signaling mediated BBB integrity under healthy circumstances.

### 2.5. Constitutive *Lef1* loss of function mutagenesis in BECs has no effects on BBB permeability

AAVs induced KO allows temporal controlled induction in adults and cell type specificity. It also elevated potential confounding effects of other inducible systems such as use of tamoxifen. Nevertheless, viral vectors have the potential to induce an immune response that can by itself influence BBB integrity^43^, and KO efficiency is somewhat lower than expected with a genetic approach. Thus, in order to validate our results, we constructed a constitutive endothelial *Lef1*-KO model by crossing *Lef1*^FL^ mice with B6.Cg-Tg(Tie2-Cre)1Ywa/J mice (Supplementary figure 5A). This mouse model is expected to provide uniform Cre expression in endothelial cells already during embryogenesis and consistently in adulthood^44^. Such approach was used in several studies to perturb Wnt components with pronounced effects on the BBB^13,15,45,46^. Genotyping and immune-fluorescent staining of cerebral cortices of 11 weeks-old mice confirmed highly efficient *Lef1*-KO resulting in a 93.95% reduction of LEF^+^ endothelial cells (from 98.86 ± 4.16% in control mice to 5.98 ± 7.51 in *Lef1*-KO mice) (Supplementary figure 5B and 5C).

The loss of *Lef1* did not impact survival during embryonic development or postnatally, and litters aligned close to the expected mendelian ratios. This is with contrast to the E12.5 lethality reported for the Tie2-cre\*Ctnnb1^fl/fl^* model^45^. Out of 22 pups 7 (31.8%) were *Lef1*^F/F^-Tie2-Cre^+^ in line with the expected 25%, all of which also survived until the age of 11 weeks when they were sacrificed for further analysis. In fact, out of the 22 pups only one did not survive until the age of 11 weeks, but since it was lacking the Tie2-Cre gene, lack of *Lef1* is unlikely to be the cause (Figure 5A).

**Figure 5.**
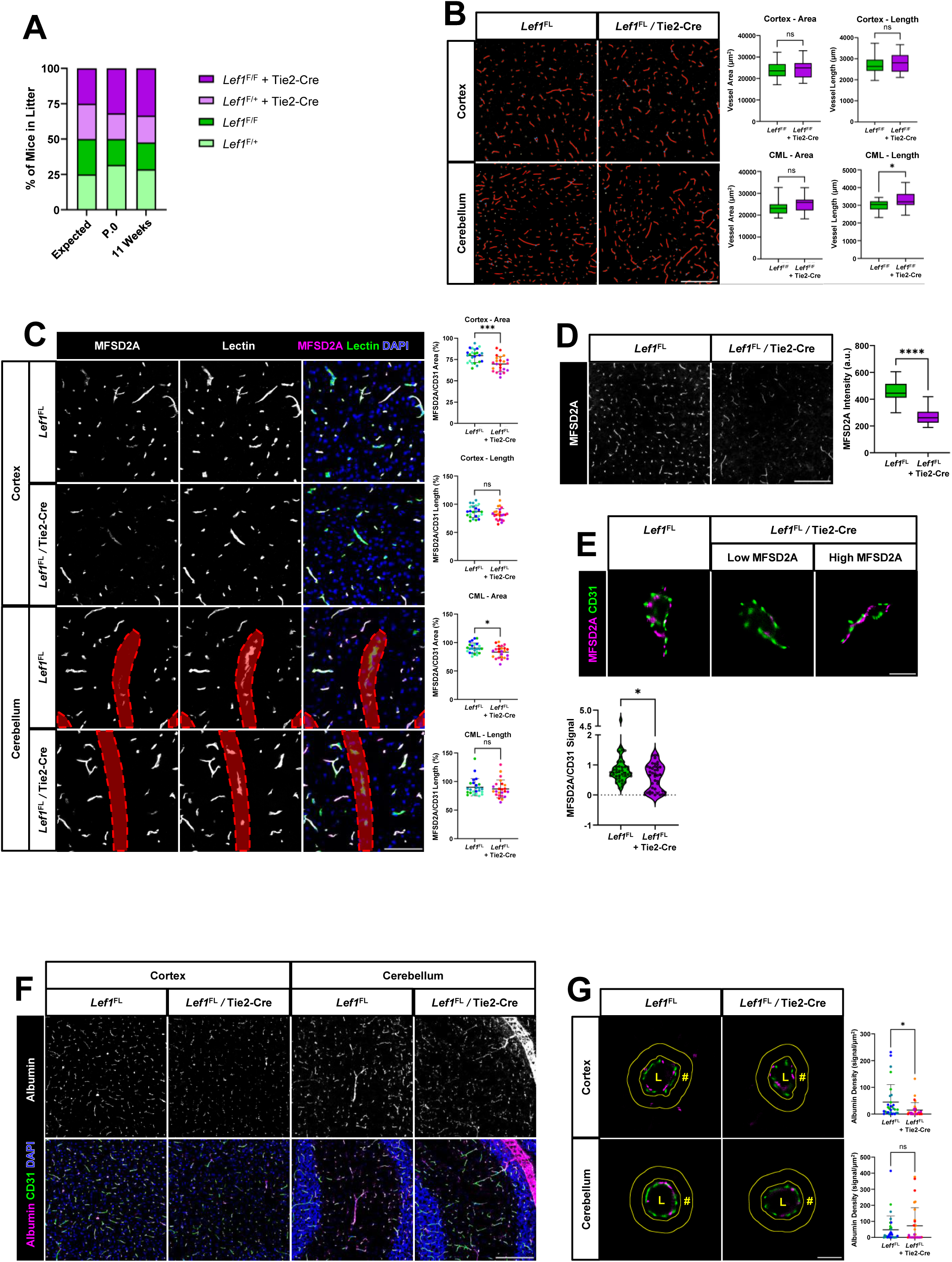
Constitutive *Lef1* loss of function mutagenesis in BECs has no effects on BBB permeability. **(A)** Fraction of the genotypes of n=22 mice used in the experiment at birth and at the age of sacrifice (11 weeks) relative to the expected ratio show no bias against Lef1-KO mice (Lef1^F/F^ Cre^+^) at birth, nor increased mortality post-natal. **(B)** Representative epi-fluorescent images of the vascular network in the cortex and the CML of *Lef1*^FL^ control mice and *Lef1*^FL^ + Tei2-Cre KO mice, show only a slight but statistically significant increase in the length of the vascular network in the CML (8.41%). Quantification was done on n=4 fields from n=6 mice, scale bar is 200µm. **(C)** Representative confocal images of MFSD2A^+^ endothelial cells immunostaining (Isolectin) in the cortex and CML of the same mice reveal a statistically significant decrease in blood vessel area in the cortex and the CML (12.30% and 7.87% respectively). Quantification of n=6 fields from n=4 mice, non-molecular layer areas in red were excluded from the quantification, scale bar is 100µm. **(D)** Representative epi-fluorescent images of MFSD2A immunostaining in the cortex show marked reduction of 37.88% in MFSD2A intensity in the *Lef1*-KO mice. Vascular MFSD2A intensity was normalized to vascular area determined by CD31 staining, and then background fluorescents was substracted. Quantification of n=6 fields from n=4 mice, scale bar is 200µm. **(E)** Super-resolution microscopy (dSTORM) of MFSD2A signals in capillaries (anti-CD31) of the cortex shows significant decrease in the average MFSD2A signal (normalized to CD31 signal). This decrease appears to represent a sub population of blood vessels of *Lef1*-KO mice having low MFSD2A levels. 10 capillaries from 4 mice from each treatment were used for quantification, scale bar is 2µm. **(F)** Representative confocal images of endogenous albumin immunostaining, in the cortex and cerebellum, **(G)** and super-resolution imaging quantification of capillaries (anti-CD31) show no significant increase in albumin leakage following *Lef1*^FL^-KO. Scale bar is 200µm for confocal and 2µm for super resolution. For quantification n=10 capillaries from n=3 mice were used, L = lumen, # = area abluminal to the endothelium quantified. Box and violin plots present data as median while the rest present the mean and standard deviation, different colors represent different mice, two-tailed Welch’s t test was used, ns = non-significant, *p < 0.05, ***p < 0.001, ****p < 0.0001.

Tie2-Cre is expected to drive *Lef1*-KO already during early stages of embryonic brain angiogenesis, and Wnt signaling is vital for these processes^13–17,44^. Therefore, we first tested for changes in vascular network properties of adult *Lef1*-KO mice brains. We did not detect any decrease in the extent of vascular network coverage in cortex or CML of *Lef1*-KO mice, but we found a slight increase of 8.41% in the average length of blood vessels in the CML (from 3003.45 ± 295.76µm per field imaged in control mice to 3255.95 ± 471.20µm in *Lef1*-KO mice) (Figure 5B).

Next, we quantified coverage of Wnt-signaling down-stream targets MFSD2A, CLDN5, GLUT1, and PLVAP in this model with confocal imaging. While CLDN5 was not significantly impacted by the loss of *Lef1* (Supplementary figure 5D), GLUT1 showed a modest decrease of 9.47% in area covered in the CML (from 76.65 ± 10.00% in the control to 69.39 ± 9.52% in the KO) (Supplementary figure 5F). PLVAP, expected increase in the absence of Wnt signaling, did not change in the *Lef1*-KO model and was nearly completely absent in blood vessels of the cortex and the CML compared to the ChP where ∼50% of the vessel area and ∼40% of the vessel length was PLVAP positive (Supplementary figure 5E). MFSD2A expression levels were more consistently impacted in the different brain areas tested, resulting in a statistically significant decrease in the area of MFSD2A^+^ blood vessels in the cortex and the CML (12.30% and 7.87% respectively) (Figure 5C). Although these changes seemed minor, they reflect a much more significant reduction in the average staining intensity of blood vessels imaged by widefield microscopy, resulting in a reduction of 37.88% (from 483.76 total pixel intensity (normalized to vascular area) in the control to 300.53pixel intensity in the KO) in MFSD2A^+^ staining intensity of cortex blood vessels (Figure 5D). dSTROM analysis further confirmed a significant decrease of 34.6% in average MFSD2A signal when normalized to CD31 signal, from an average MFSD2A to CD31 signal ratio of 0.900 ± 0.697 in the control mice to 0.589 ± 0.438 in *Lef1*-KO mice (Figure 5E). Taken together, the effect of *Lef1*-KO under the Tie2-Cre control seems to have a somewhat stronger effect on some of Wnt-signaling related BBB genes at the protein levels.

Finally, reduced MFSD2A expression is related to increase permeability mediated by lowering inhibition of transcytosis. Such a mechanism was demonstrated for the CML hyper-permeability upon ECs *Ctnnb1* loss^21^. In contrast, constitutive endothelial *Lef1*-KO starting during embryonic development did not impact BBB permeability in 11 weeks old mice. We immunostained for albumin, and quantified albumin signal intensity up to 1µm from blood vessels in the cortex and CML. Yet, we did not find any increase in BBB permeability in *Lef1*-KO mice either. In fact, a statistically significant decrease in extravascular albumin signal was observed in the cortex of *Lef1*-KO (from 44.94 ± 65.35% in the control to 15.00 ± 28.17% in the KO) (Figure 5F and 5G). This leads us to conclude that *Lef1* is not necessary for BBB functionality in adult mice.

## 3. Discussion

Over the years comprehensive research has been conducted showing the importance of the canonical Wnt signaling pathway for the formation of the BBB as well as for its maintenance during adulthood^13,16,18–22,25^. More specifically, recent studies using 8-10 weeks old *Ctnnb1^fl/fl^; Cdh5-Cre^ER^* mice show significant reduction in Wnt signaling, severe dysregulation of barrier related gene expression (e.g. tight-junction proteins, Cav1, and Mfsd2a), and increased BBB permeability only 3 days after the last tamoxifen injection (8-10 days after the first injection). This is reported to be followed by complex neurological pathologies mere 1-2 week after tamoxifen injection pointing towards the critical role of β-catenin in the maintenance of the BBB^20,21^.

In our study we focused on the role of the *Tcf/Lef* transcription factors, particularly *Lef1*, in BBB maintenance. This family of transcription factors has long been established as the main partners of β-catenin in the transcriptional regulation of the genes downstream of the canonical Wnt signaling pathway^47^, yet their role in forming and maintaining a functional BBB has not been directly explored. We therefore began our inquiry into the role of *Tcf/Lef* in BBB maintenance by exploring the effect of BEC specific *Lef1*-KO in adult mice. We demonstrate that while the loss of *Lef1* has a significant impact on Wnt signaling, the effect on BBB specific gene expression is modest, and damage to BBB integrity is not detectable even 4 weeks following *Lef1-*KO induction.

The *Tcf/Lef* family contains 4 transcription factors that all form complexes with β-catenin, of which *Lef1* acts mostly as an activator, *Tcf7l1* generally acts as a repressor, and *Tcf7* and *Tcf7l2* as either activators or repressors of Wnt target genes. The role of *Tcf/Lef* as either activator or suppressor is highly dependable on the cellular context and differs between organisms, developmental stages, and tissues where they are expressed. Additionally, there tends to be overlapping expression patterns of the different *Tcf/Lef*, making it common that two or more are co-expressed in the same cell and create functional redundancy^29^. With that said, examples of single KO of one *Tcf/Lef* gene, particularly *Lef1*, resulting in observable effects also exist^48,49^. This, together with previously published single cell RNA data showing much elevated *Lef1* expression in the brain endothelium relative to the non-barrier endothelium^32,34^, led us to explore if *Lef1* might be necessary for the β-catenin mediated BBB maintenance.

Based on our results, we conclude that while *Lef1* does play a role in BEC Wnt signaling, it is not necessary for BBB maintenance and formation, since even using *Lef1^fl/fl^; Tie2-Cre* in which endothelial *Lef1*-KO occurs already during embryonic development^44^, no significant impact on BBB integrity could be observed in adult mice. We can postulate three possible explanations for this discrepancy, beyond the possibility of compansation: (1) *Lef1* effect on Wnt signaling (detected by *Axin2*, *Nkd1*, *Apcdd1*, *Spock2*, and *Notum* reduction) reflects a much-reduced effect compared to the one resulting from *Ctnnb1*-KO and is thus might be insufficient to induce significant changes to BBB functionality. (2) *Lef1* might be obligatory for the regulation of BBB nonspecific genes like *Axin2* and *Nkd1*, but regulation of barrier specific genes like *Mfsd2a* and *Glut1* might also be regulated through different transcription factors. (3) The BBB disruption resulting from *Ctnnb1*-KO or by interfering with Wnt signaling upstream of β-catenin might impact directly cell junction integrity, and not necessarily transcriptional regulation by β-catenin.

As mentioned before, the effects of *Lef1*-KO on the Wnt signaling targets *Axin2*, *Nkd1*, *Apcdd1*, *Spock2*, *Notum*, and *Lef1* that we observed was less consistent with the reported effects resulting from *Ctnnb1*-KO^20^,with *Axin2* changes not being statistically significant, and total *Lef1* RNA levels not being impacted at all in *Lef1*-KO BECs. While the differences in results could be explained by differences in experimental procedures, they could also indicate a partial effect of *Lef1*-KO compared to *Ctnnb1*-KO. This reduced effect on Wnt signaling might then cause an even smaller effect on BBB-specific gene expression which in turn is inadequate to impact BBB functionality.

On the other hand, if the observed impact of *Lef1*-KO on Wnt signaling targets *Axin2*, *Nkd1*, *Apcdd1*, *Spock2*, and *Notum* reflects a substantial effect comparable to the one observed in *Ctnnb1*-KO models, while the impact of BBB integrity remains minimal, then this could hint at differences in the transcriptional regulation of barrier specific and barrier non-specific Wnt signaling targets. *Axin2*, *Nkd1*, *Apcdd1*, *Spock2*, and *Notum* though well-established Wnt signaling target genes don’t seem to have any direct effect on BBB functionality. *Axin2*, *Nkd1*, *Apcdd1* and *Notum*, though up-regulated by β-catenin, do themselves repress Wnt signaling by various mechanisms effectively creating a negative feedback loop for Wnt signaling^50–53^. *Spock2* meanwhile, is a member of the SPARC family that are comprised of extracellular matrix associated proteins^54^. Therefore, it is not necessary that significantly effecting those general Wnt signaling target genes by the means of Lef1-KO will also lead to a comparable effect on BBB specific Wnt signaling target genes. Transcriptional regulation of barrier critical genes involving β-catenin might be more robustly compensated or even entirely regulated by other *Tcf* family members, particularly by *Tcf7* as indicated by accessible TCF7 and LEF1 binding sites upstream of *Slc2a1* and *Mfsd2a* in BECs^30,31^.

Lastly, β-catenin is not only functioning as a transcription factor but plays also a direct role in BBB integrity by being a key part of the adherence junction complex. In this context β-catenin together with α-catenin and p120 create a complex that links VE-cadherin to the actin cytoskeleton^55^. While some evidence suggests that plakoglobin can compensate for β-catenin loss in the adherence junctions^19,45^, others show that *Ctnnb1*-KO leads to disruption of adherence junctions^20^. Therefore, if *Ctnnb1*-KO directly effects BBB integrity by opening the adherens junctions, further BBB breakdown might possibly follow without the need for direct transcriptional dysregulation of BBB specific Wnt signaling targets^55^.

In light of our results, our leading hypothesis to the effect of BEC specific *Lef1*-KO is that one or more of the other *Tcf* members are compensating for the lack of *Lef1*. Probably the much more modestly elevated levels of *Tcf7* in BECs act to allow redundancy for *Lef1* activity, at least under homeostatic conditions. In the current study, we did not address the option that lack of endothelial *Lef1* expression might have an exacerbating effect on BBB damage during stress or insult like in the case of ischemic strokes. In order to further elucidate the vital role that the canonical Wnt signaling pathway plays in BBB maintenance, in future research we plan to address the possibility of Tcf7 compensation by testing the effect of a *Lef1*-*Tcf7* double KO in adult mice. These insights will allow for further advancements of therapeutic strategies to further BBB recovery following damage, or to manipulate the BBB for drug delivery into the CNS.

### 4. Material and Methods

### 4.1. Animals and treatments

A stock of *Lef1*^FL^ (B6.Cg-*Lef1^tm1Hhx^*/J mice, JAX stock #030908, originally produced by Dr. Hai-Hui Xue)^35^ was kindly provided by the lab of Prof. Mona Dvir-Ginzberg (Institute of Biomedical and Oral Research, Hebrew University of Jerusalem, Israel). For some experiments *Lef1*^FL^ were crossbreed into stock Tie2-Cre mice (B6.Cg-Tg(Tek-cre)1Ywa/J, JAX stock #008863)^44^. For AAV titer and injection time calibration C57BL/6JOlaHsd mice were used. All animals were housed in SPF conditions and treated according to institutional guidelines approved by the Institutional Animal Care and Use Committee (IACUC) at Hebrew University. *Lef1*^FL^ genotype was confirmed using the primers detailed in table 1.

**Table 1.**
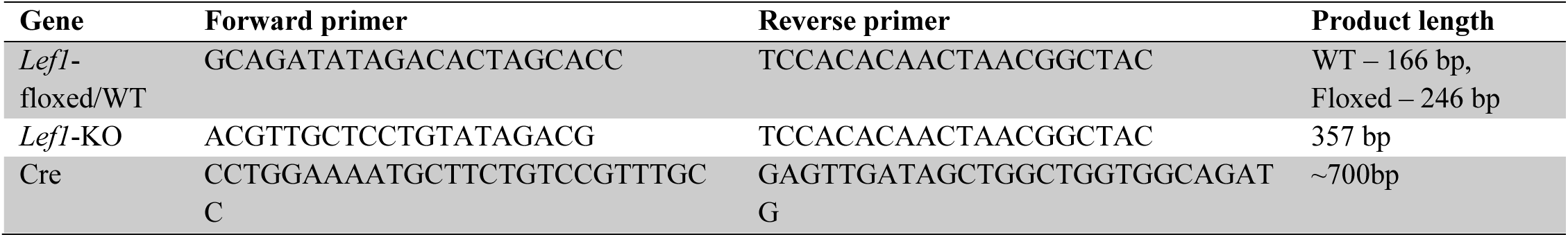
Primers for genotyping.

Adult mice were intravenously injected with 100µl, and a total titer of 1-2e11vg of kiCAP-AAV-BI30 capsid (Addgene #183749)^36^, containing either pAAV-CAG-Cre-3xmiR122-WPRE-HGHpA (Addgene #183776)^36^ or pAAV-CAG-SV40NLSf-GFP-3xmiR122-WPRE-HGHpA (Addgene #183775)^36^ packaged by ELSC Viral Core Facility were used. The mice were then kept for 3-4 weeks in the facility, until their sacrifice.

### 4.2. Capillary isolation and RNA extraction

Deeply anesthetized 13 weeks old Lef1^FL^ mice injected with AAV-BI30 (as described above) were perfused for 2-3 minutes with ice-cold PBS. The brain was then extracted, cleaned of the meninges, and the cortex was dissected and placed into 2mL cold MVB solution (15 mM HEPES, 147 mM NaCl, 4 mM KCl, 3 mM CaCl_2_+2H_2_O, 1.2 mM MgCl_2_+6H_2_O, in double distilled water (DDW), titrated to pH 7.4, and filtered for stock solution) supplemented with 5 mM D-glucose and 0.5% bovine serum albumin (BSA) added to stock solution immediately prior to the experiment. Cortices from each mouse were then separately homogenized using a power homogenizer (WHEATON, #903475, 20 stokes at speed 3), the lysate was transferred to a 15 mL tube, and the glass tube used for homogenization was washed with an additional 2 mL MVB. Lysate was then centrifuged in a bucked centrifuge cooled to 4 °C at 400 g for 10 minutes, the supernatant was removed, and the pellet was resuspended in 4 mL 25% BSA in PBS, vortexed thoroughly, and centrifuged at 2000 g for 30 minutes. Top myelin layer and supernatant were carefully removed, and the pellet resuspended in 1 mL MVB. Next the pellet was filtered through a 100 µm mesh to remove large debris, and then the flow through was filtered again through a 40 µm. Capillaries, captured on the 40 µm, were then immediately lysed using 800 µL RNA lysis buffer from the Quick-RNA MicroPrep kit (Zymo Research, #R1051).

The RNA was then extracted using the MicroPrep kit as per manufacturer’s instructions. The 15 µL eluted RNA was then turned into cDNA using the High Capacity cDNA Reverse Transcription Kit (Applied Biosystems, #43-749-66) as per manufacturer’s instructions. Resulting cDNA was then diluted with 80 µL nuclease free water and stored at -20 °C until further analysis.

### 4.3. qPCR

384 well PCR plates (BIO-RAD, #HSP3805) where loaded cDNA and primers as detailed in table 2 where loaded as triplicates. Each reaction contained 1 µL cDNA, 3.8 µL nuclease free water, 0.15 µL primers (150nM), and 5 µL PowerSYBR Green (Applied Biosystems, #4367659). The plate was then sealed (BIO-RAD, #MSB1001) and spun down at 700 g for 1 minute and then run in C1000 Touch Thermal Cycler (BIO-RAD, # 1851138) with the CFX384 Real-Time System (BIO-RAD, # 1845385) and running Bio-Rad CFX Maestro 2.3 with the following protocol: 95 °C for 10 minutes, 40 cycles of 95 °C for 15 and 60 °C for 1 minute, closing with 65 °C for 15 seconds. Threshold cycle (C_t_) was normalized to *Gapdh* and to the average C_t_ of the control samples before converting to 2ΔΔC_t_.

**Table 2.**
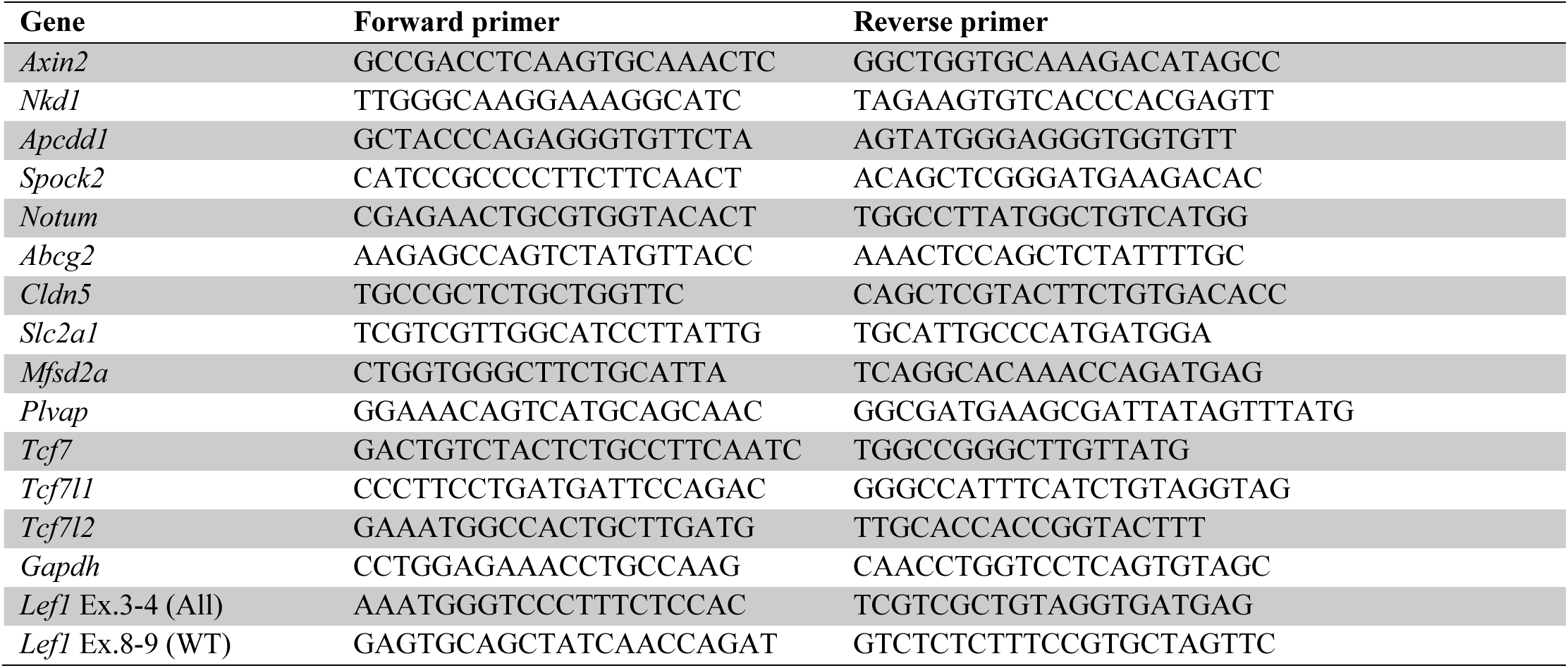
qPCR primers.

### 4.4. Tracer Injection

Deeply anesthetized 14 weeks old Lef1^FL^ mice injected with AAV-BI30 (as described above) were perfused for 5 minutes with sulfo-NHS-biotin (4 mg/20 g mouse body weight, Thermo Scientific, #21217, dissolved in 20 ml phosphate buffered saline (PBS)). Brains were dissected and fixed in 4% paraformaldehyde (PFA) as described under tissue preparation section.

### 4.5. Tissue preparation

For fixed tissue samples mice were euthanized by cervical dislocation under isoflurane anesthesia. The brain was then extracted into 4% PFA in PBS and kept at 4 °C overnight. Samples were then thoroughly washed in PBS before transferred into 30% sucrose solution. Samples were then kept at 4 °C until the tissue sunk to the bottom of the vile at which point they were frozen in O.C.T compound (Scigen, #4586).

For fresh-frozen tissue samples mice were anesthetized using a Ketamine Xylazine mix injected intraperitoneal. They were then perfused using 1% PFA in PBS with heparin (10-20 units/ml) for 5 minutes, before the brain was extracted and fresh-frozen in O.C.T compound.

All samples were stored at -80 °C, and 6-10 µm sections created using a cryostat (Leica, #CM1950).

### 4.6. Immunohistochemistry

For fresh frozen slides, post fixation was performed prior to staining using 4% PFA for 10 minutes at room temperature followed by 3 washes with PBS.

All slides were treated with blocking solution (10% BSA, 10% horse serum, 0.5% triton-x100, in PBS) for 2 hours at room temperature prior to staining. Slides were then incubated with primary antibodies (see antibodies table for details; Table 3) in incubation buffer (2.5% BSA, 2.5% horse serum, 0.5% triton-x100, in PBS) over night at 4 °C and then washed off with PBS 3 times. Secondary antibodies (see fluorophores table for details; Table 4) in incubation buffer were then applied for another 1 hour at room temperature followed by washing with PBS and mounting with DAPI-fluoromount-G (southern biotech, #0100-20).

**Table 3.**
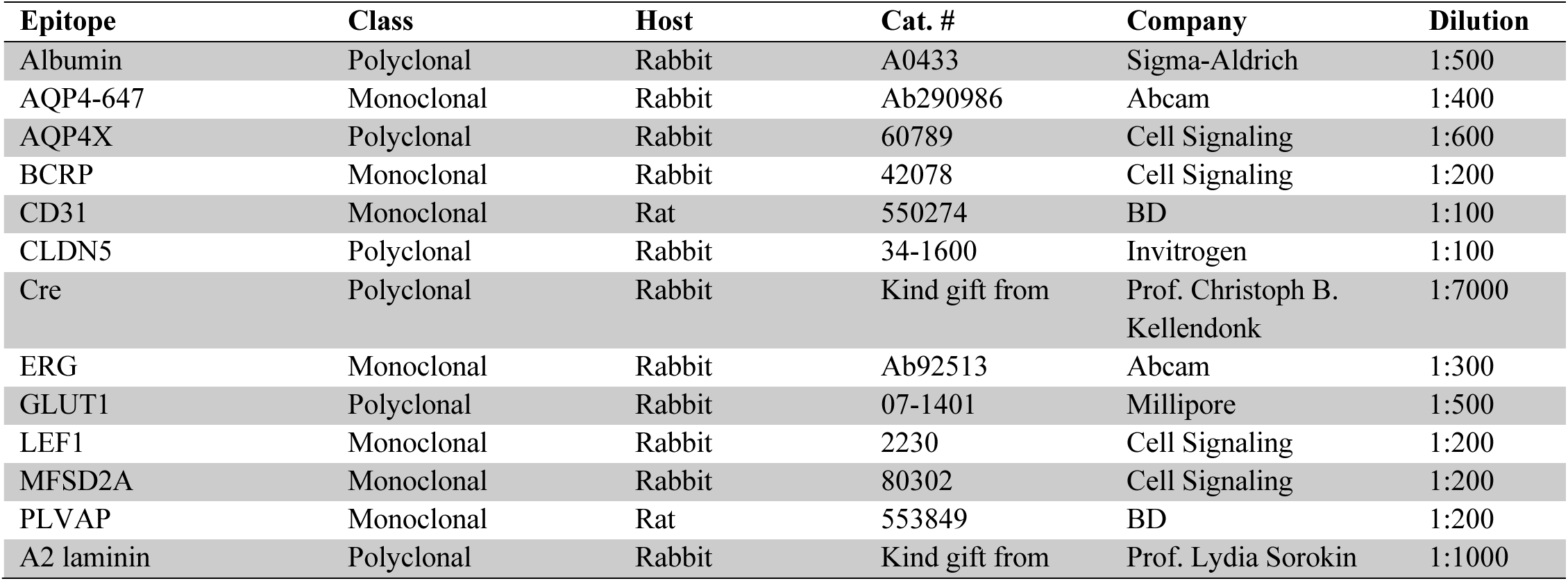
Primary antibodies used.

**Table 4.** Fluorophores used.

| Fluorophore | Isotype | Cat. # | Company | Dilution |
| --- | --- | --- | --- | --- |
| Alexa fluor 647 | Anti-rat IgG | 712-605-153 | Jackson | 1:500 |
| Alexa fluor 488 | Anti-rabbit IgG | 711-545-452 | Jackson | 1:200 |
| Cy3 | Anti-rabbit IgG | 711-165-152 | Jackson | 1:300 |
| Alexa fluor 488 | Anti-rat IgG | 712-545-153 | Jackson | 1:200 |
| Isolectin GS-IB<sub>4</sub> Alexa fluor 647 | α-D-galactosyl | I32450 | Invitrogen | 1:300 |
| Streptavidin Alexa fluor 647 | Biotin | S32357 | Molecular Probes | 1:1000 |
| Alexa fluor 647 | Anti-rabbit IgG | 711-605-152 | Jackson | 1:500 |
| CF568 | Anti-rabbit IgG | 20098 | Biotum | 1:400 |

For immuno-staining with isolectin, first a stock solution of 1 µg/µL isolectin GS-IB4 Alexa Fluor 647 conjugate (Invitrogen, #I32450) in 0.1 M CaCl_2_ in DDW pH 7.3. This solution was applied at 1:300 together with the secondary antibodies in PB-lec (1% triton-x100, 0.1 mM CaCl_2_, 0.1 mM MgCl_2_, 0.1 mM MnCl_2,_ in PBS) instead of the incubation buffer mentioned above.

For double staining with anti-rabbit antibodies (for anti-LEF1/anti-Cre with anti-ERG), first staining was performed with either anti-LEF1 or anti-Cre antibodies in the manner described above but with 1.5 hours incubation of secondary antibody instead of 1 hour in order to saturate anti-rabbit binding sites. Then the samples were washed thoroughly with PBS and stained for 2 hours at room temperature with anti-ERG antibodies, then washed, incubated for 1 hour with the second secondary antibody, washed again and mounted.

### 4.7. Fluorescence microscopy

Confocal mages were captured using Nikon A1R+ confocal microscope, objective X20 and X40, and Nis-Elements software.

Epi-fluorescence images were taken using an Olympus BX51, 10x/0.30, 20x/0.50, and 40x/0.75, with Andor Zyla camera, and Nikon NIS elements software (version D4.5).

### 4.8. Fluorescence microscopy quantification

For analysis and quantification of confocal end epi-fluorescence microscopy data we utilized Fiji based on ImageJ^56^. For quantifying protein coverage of capillaries and vascular brain perfusion (see Figure 5B) we also used AngioTool64 0.6a^57^.

### 4.9. dSTORM imaging

We used a dSTORM (direct stochastic optical recon-struction microscopy) system, which allows imaging at approximately 20 nm resolution by using photo-switch-able fluorophores (all dSTORM imaging was done on TIRF mode). 8μm brain slices were mounted on poly-D-lysine coated coverslips (no. 1.5 H, Marienfeld-superior, Lauda-Königshofen, Germany). dSTORM imaging was performed in a freshly prepared imaging buffer containing 50 mM Tris (pH 8.0), 10 mM NaCl and 10% (w/v) glucose with an oxygen-scavenging GLOX solution (0.5 mg/ml glucose oxidase (Sigma-Aldrich)), 40 μg/ml catalase (Sigma-Aldrich), 10 mM cysteamine MEA (Sigma-Aldrich), and 1% β mercaptoethanol ^58–60^. A Nikon Ti-E inverted microscope was used. The N-STORM (Nikon STORM system) was built on TIRF illumination using a 1.49 NA X100 oil immersion objective and an ANDOR DU-897 camera. 488, 568 and 647 nm laser lines were used for activation with cycle repeat of ∼ 4000 cycles for each channel. Nikon NIS Element software was used for acquisition and analysis; analysis was also performed by ThunderSTORM (NIH ImageJ^61^).

### 4.10. dSTROM quantification

The dSTORM approach we used is based on labeling the target protein with a primary antibody and then using a secondary antibody conjugated to a fluorophore. Thus, resolved signals represent a location that is approximately 40 nm from the actual epitope (assuming the approximation of the two antibodies’ length in a linear conformation). The number of signals represents an amplification of the actual target numbers. Amplification corresponds to the primary antibody in the case of a polyclonal antibody (assuming binding to several epitopes in the same protein, which could be reduced by the use of monoclonal antibodies). Amplification also corresponds to several secondary antibodies binding to a single primary antibody and to several fluorophores attached to a single secondary antibody. Nevertheless, resolution of approximately 20 nm allows us to separate signals and to use these as proxies to the abundance of target molecules, which can reliably be used to compare different states^9,62^.

Single molecule localization microscopy (SMLM) results in point patterns having specific coordinates of individual detected molecules. These coordinates are typically summarized in a ‘molecular list’ (provided by ThunderSTORM analysis (NIH ImageJ)^61^.

### 4.11. Statistical analysis

All comparisons were performed by two-tailed Welch’s t test, (as indicated in the figure legends), assuming normal distribution for all but the qPCR data for which a lognormal distribution was assumed, p < 0.05 was considered significant. GraphPad Prism 10.6.1 for Windows (GraphPad Software, San diego, California, USA) was used for all graphs except for figure 1A, for which R 4.3.2 in conjunction with RStudio 2024.12.0+467 were used.

## 5. Acknowledgments

We would like to thank; the Ben-Zvi group for scientific and writing inputs, Dr. Hai-Hui Xue for permission to use the Lef1^FL^ mice, Dr. Mona Dvir-Ginzberg for supplying us with the *Lef1*^FL^ mice, Dr. Vanlandewijck and Dr. He for permission and assistance in using their single cell RNA sequencing data, the ELSC Viral Core Facility for providing us with the AAVs, Prof. Lydia Sorokin for the anti-Laminin-a2 antibodies, and Prof. Christoph B. Kellendonk for the anti-Cre antibodies.

## Supplementary Figure Legends

**Figure S1.**
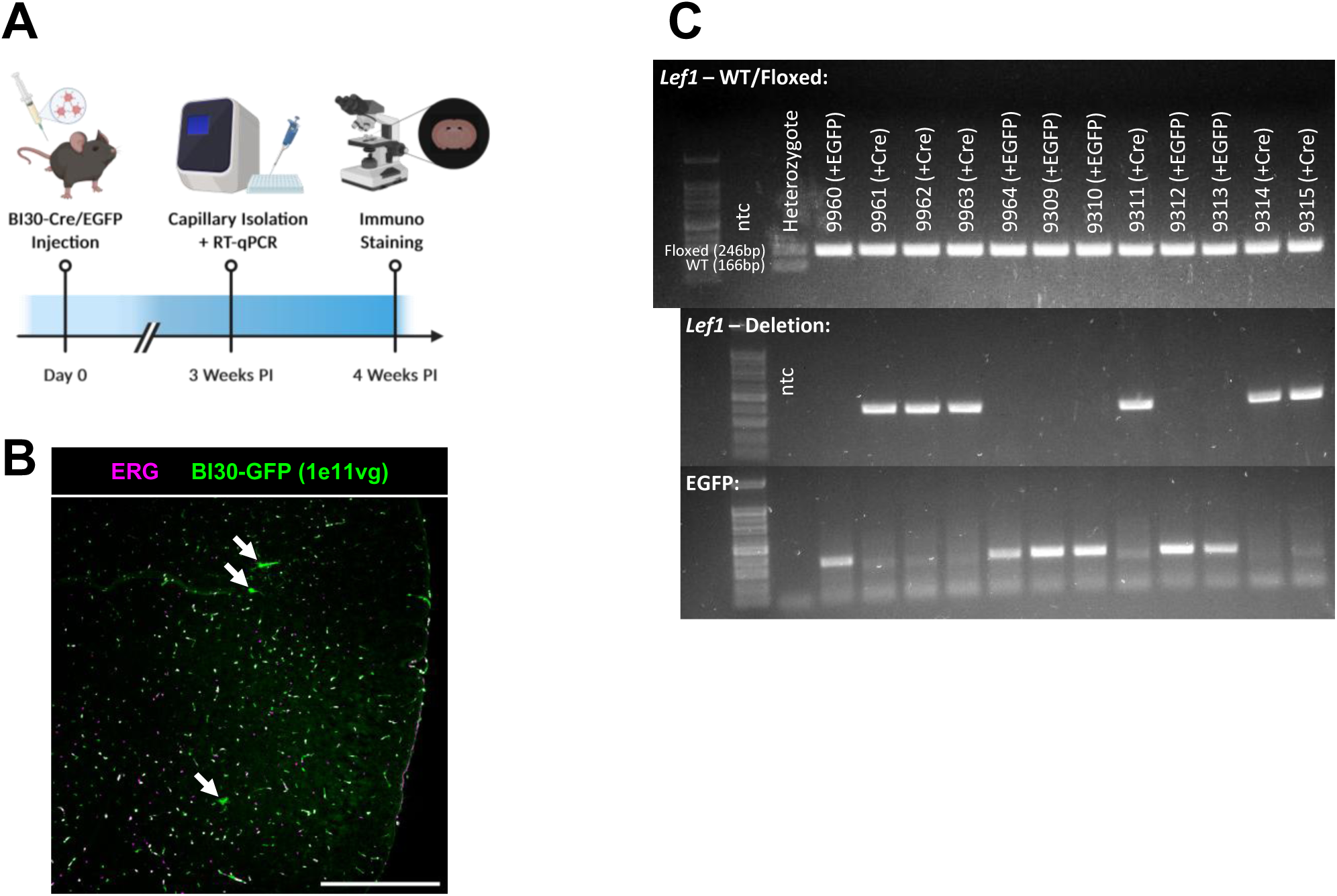
Use of endothelial tropic AAVs to induce *Lef1* loss of function mutagenesis in adult mice. **(A)** Schematic illustration of the experimental design: AAV-BI30-Cre/EGFP is injected retro-orbital at 11-14 weeks, tissues are then taken for qPCR 3 weeks post injection (PI) and for immune-staining 4 weeks PI. Created in BioRender. Yeretz peretz, Y. (2025) https://BioRender.com/4v5gdex **(B)** Representative epi-fluorescent image of the cerebral cortex of a C57BL/6JOlaHsd mouse injected with communally used AAV-BI30-EGFP titers, demonstrating mostly vascular infection (endothelial nuclei are marked with ERG) with only minimal non-endothelial infection (white arrows). Scale bar is 400µm. **(C)** Representative genotyping result of the litter used for the qPCR experiments. Primers for the floxed or WT *Lef1* allele, for *Lef1* after excision of exons 7-8, and for Cre were used.

**Figure S2.**
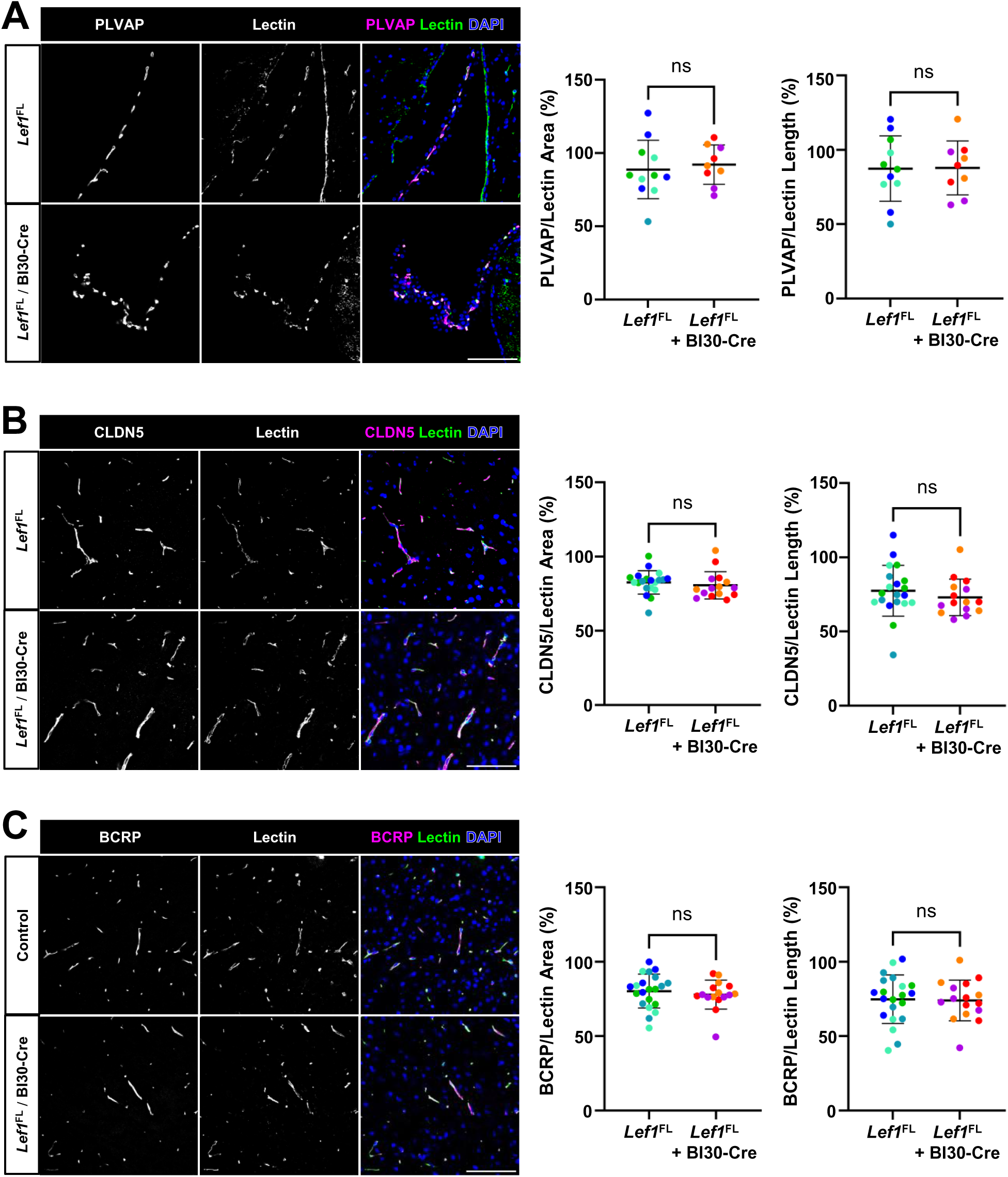
BECs *Lef1* loss of function has no detectable effect on canonical Wnt signaling induced expression of BBB related downstream targets. **(A)** Representative confocal images of immune-staining PLVAP^+^ in the ChP of AAV-BI30-Cre injected *Lef1*^FL^-KO mice or un-injected controls, show robust PLVAP coverage in non-barrier blood vessels (Isolectin). Confocal images of **(B)** CLDN5^+^ **(C)** and BCRP^+^ immune-staining show no reduction in the coverage of blood vessels in the cortex (stained with Isolectin) of *Lef1*-KO mice compared to control mice. All quantification was done on n=5 fields of n=4 control and n=3 *Lef1*-KO mice, data is presented as mean and standard deviation, different colors represent individual mice, two-tailed Welch’s t test was used, ns = non-significant, scale bars are 100µm.

**Figure S3.**
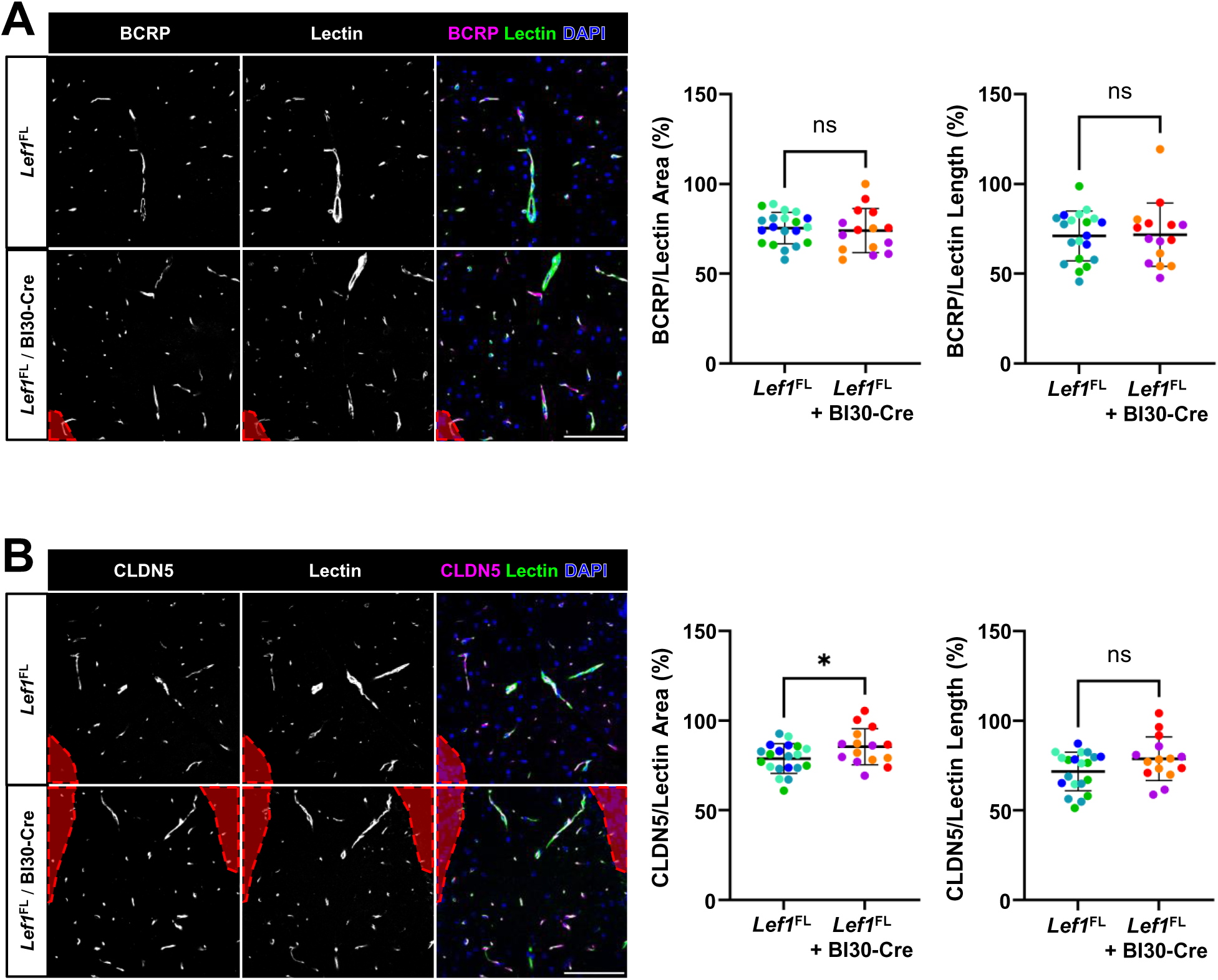
Loss of *Lef1* function in BECs has no observable effect on the expression of downstream Wnt signaling targets in susceptible brain areas. **(A)** Representative confocal images of BCRP^+^ and **(B)** CLDN5^+^ endothelial cells immune-staining, of the CML (Isolectin) show no reduction in coverage, rather a slight increase of 8.34% in CLDN5^+^ vascular area. Non-molecular layer areas (namely the granular layer of the cerebellum) in red were excluded from the quantification. n=5 fields of n=4 control and n=3 *Lef1*-KO mice were used, data is presented as mean and standard deviation, different colors represent individual mice, two-tailed Welch’s t test was used, ns = non-significant, *p < 0.05, scale bars are 100µm.

**Figure S4.**
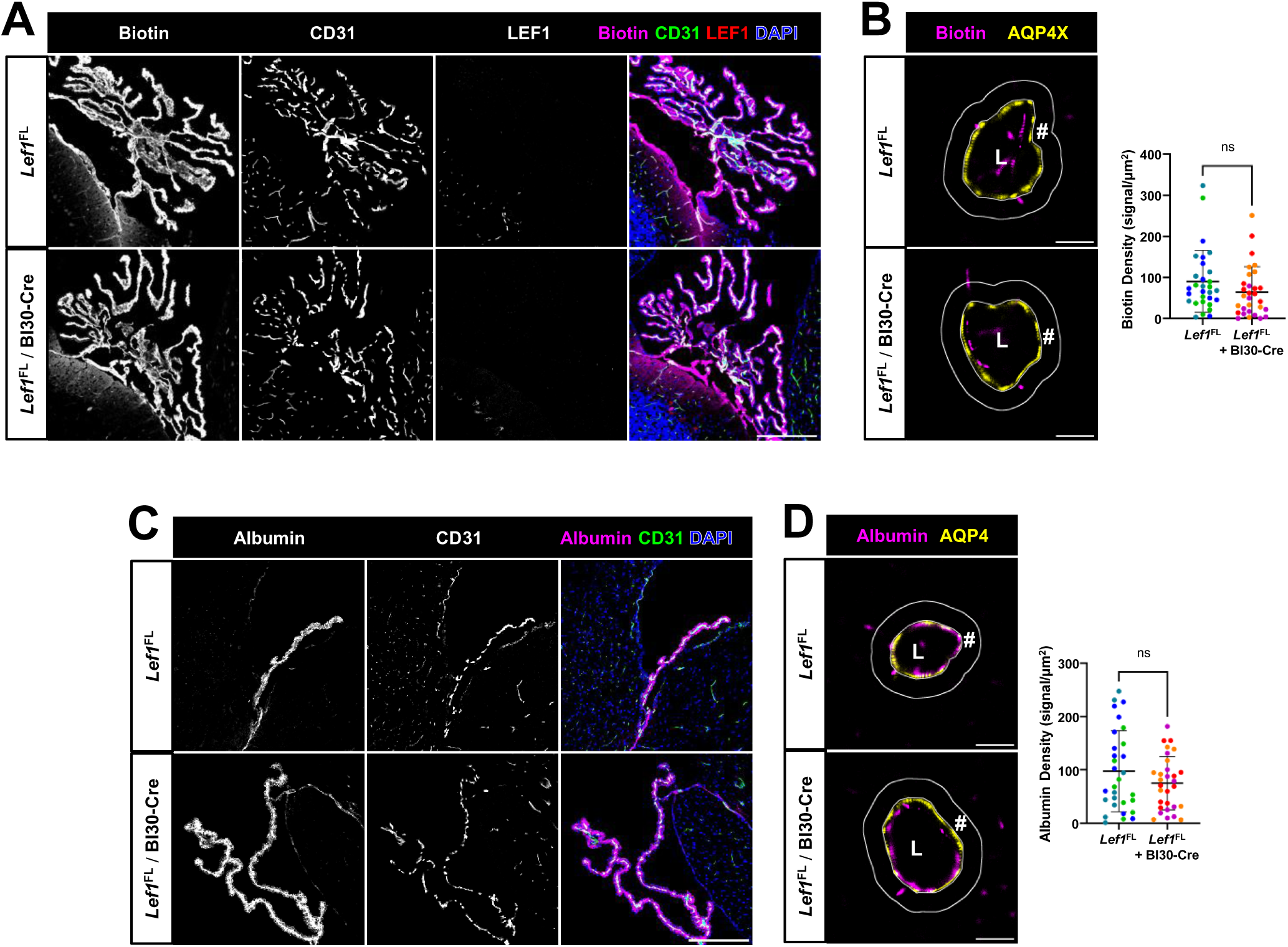
Loss of *Lef1* function in BECs has no effects on BBB permeability. **(A)** Confocal images of Biotin challenges in the ChP of the 4^th^ ventricle show considerable biotin leakage in the ChP and around the attachment region**. (B)** Quantification of super-resolution (dSTORM) images of capillaries from the cortex of AAV-BI30- Cre injected *Lef1*^FL^-KO mice or un-injected controls show no significant increase in biotin signal 1 µm from the astrocytic and-feet (anti-AQP4X). Confocal images of endogenous albumin immunostaining in the ChP of the lateral ventricles show considerable albumin leakage in the ChP**. (B)** Quantification of super-resolution (dSTORM) images of capillaries from the cortex of AAV-BI30-Cre injected *Lef1*^FL^-KO mice or un-injected controls show no significant increase in albumin signal 1 µm from the astrocytic and-feet (anti-AQP4). L = lumen, # = area quantified. Scale bar is 200µm for confocal and 2µm for super resolution. Quantification of super-resolution data was done on n=10 capillaries of n=3 mice of each treatment, data is presented as mean and standard deviation, different colors represent individual mice, two-tailed Welch’s t test was used, ns = non-significant.

**Figure S5.**
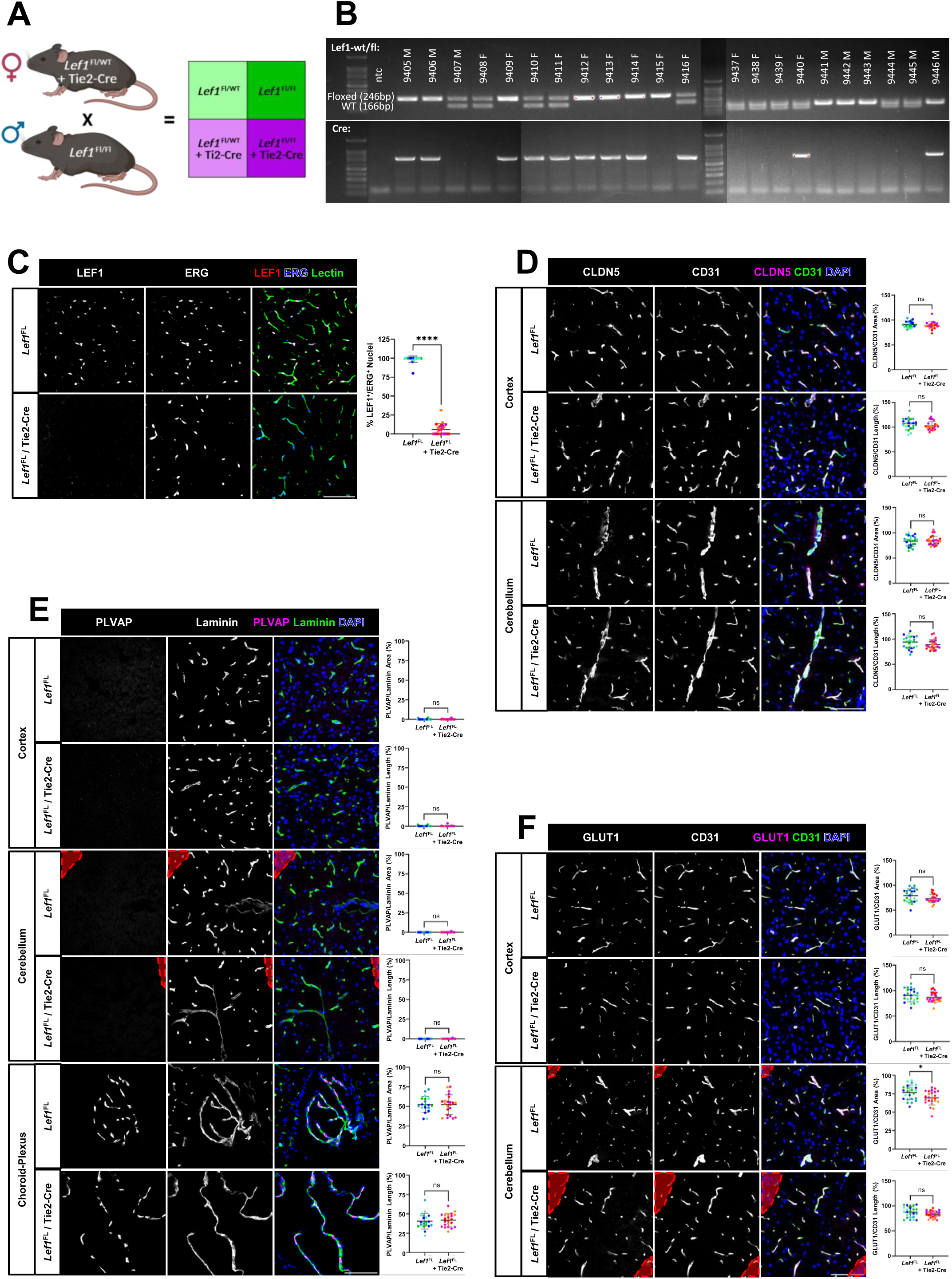
Constitutive *Lef1* loss of function mutagenesis in BECs has no effects on BBB permeability. **(A)** Schematic illustration of the cross breading performed in the experiment and expected genotypes, all four genotypes are expected at an equal rate of 25%. Created in BioRender. Yeretz peretz, Y. (2026) https://BioRender.com/wm29sqo **(B)** Genotyping result of the liter used in assessing survival rates and used in the further experiments, using primers for the floxed or WT *Lef1* allele and for Cre. **(C)** Representative epi-fluorescent images of LEF1^+^ brain endothelial cells immune-staining, in the cortex of *Lef1*^FL^ control mice and *Lef1*^FL^ + Tei2-Cre KO mice show a 93.95% reduction in LEF^+^ endothelial cells, Isolectin is used to mark endothelial cells and ERG to mark endothelial nuclei. Scale bar is 100µm. **(D)** Representative confocal images of CLDN5^+^, **(E)** PLVAP^+^, **(F)** and GLUT1^+^ endothelial cells immune-staining (anti CD31 or anti-laminin) in the cortex, CML, and ChP of the same mice, reveal a statistically significant reduction only in the area of GLUT1^+^ blood vessels (a 9.47% reduction). Red areas mark areas excluded in the quantification, namely the granular layer of the cerebellum. n=5 fields of n=4 control and *Lef1*-KO mice were used, data is presented as mean and standard deviation, different colors represent individual mice, two-tailed Welch’s t test was used, ns = non-significant, *p < 0.05, ****p < 0.0001, scale bars are 200µm.

